# Retinoid acid receptor β mechanically regulates the activity of pancreatic cancer cells

**DOI:** 10.1101/2022.08.10.503236

**Authors:** Carlos Matellan, Dariusz Lachowski, Ernesto Cortes, Stephen Thorpe, Armando E. del Río Hernández

## Abstract

Pancreatic ductal adenocarcinoma (PDAC) is the most common and lethal form of pancreatic cancer, characterised by stromal remodelling, elevated matrix stiffness and high metastatic rate. Retinoids, compounds derived from vitamin A, have a history of clinical use in cancer for their anti-proliferative and differentiation effects, and more recently have been explored as anti-stromal therapies in PDAC for their ability to induce mechanical quiescence in cancer associated fibroblasts. Here we demonstrate that retinoic acid receptor β (RAR-β) transcriptionally represses myosin light chain 2 (MLC-2) expression, a key regulatory component of the contractile actomyosin machinery. In turn, MLC-2 downregulation results in decreased cytoskeletal stiffness and traction force generation, impaired response to mechanical stimuli via mechanosensing and reduced ability to invade through the basement membrane.

## 1. Introduction

Pancreatic ductal adenocarcinoma (PDAC) constitutes the vast majority (90%) of pancreatic cancers [1] and is one of the most aggressive forms of cancer, being the 5^th^ most common cause of cancer related death in the UK [2] and the 7^th^ worldwide [3]. The 5-year survival rate has remained below 4% in the UK since the 1970s, and has barely improved globally (6-9% between 2014 and 2018) [3], in part due to the fast progression of the disease, its metastatic potential and the difficult diagnosis. This dismal prognosis and the ineffectiveness of classical treatments calls for novel therapeutic strategies to tackle the burden of pancreatic cancer.

In the last decade, the biomechanical interaction between the cancer cell and the tumour microenvironment has gained interest as a fundamental factor driving the onset and progression of cancer [4]. From a biomechanical point of view, PDAC is characterised by a highly stiff and fibrotic tumour microenvironment. This aberrant, collagen-rich tumour stroma is populated by activated pancreatic stellate cells (PSCs), which remodel the extracellular matrix (ECM) into a cancer-permissive microenvironment [5, 6]. This microenvironment, in turn, promotes epithelial-to-mesenchymal transition (EMT) [7, 8], guides cancer cell migration [9-11], increases secretion of MMPs [12, 13] and promotes chemoresistance [14], and its mechanical properties (e.g. increased stiffness) correlate with metastatic potential and response to treatment [15]. Bidirectional crosstalk between the tumour cells and the activated PSC maintains the tumour microenvironment, sustains PSC activation and drives cancer cell malignancy [16]. In addition, cancer cells appear more mechanically active than their healthy counterparts, including increased migration speed and contractility, highlighting the importance of the biomechanical milieu in cancer progression.

The retinoic acid receptors RAR-α, RAR-β and RAR-γ are the three members of the retinoic acid receptor (RAR) subfamily of nuclear receptors, which play a wide variety of roles in embryonic development, morphogenesis and proliferation [17]. RARs can be cytoplasmic or nuclear, and are activated by the binding of retinoids, the active forms of vitamin A. When activated, RARs bind their transcription partners retinoid X receptors (RXRs) to form heterodimeric transcription factors that bind specific sites of target genes known as retinoid acid response elements (RAREs) [18]. Of the three RAR family members, RAR-α is ubiquitously expressed, while RAR-β and RAR-γ are tissue specific [19]. Interestingly, the expression of RAR-β is lost or downregulated in a variety of carcinomas, including breast [20, 21], lung [22, 23], liver [24] and pancreatic [25, 26] cancer among others. This dysregulation in the expression of RAR-β often accompanies the early stages of cancer and may be concomitant with its development [27].

Retinoids have been explored as a treatment for cancer in diverse contexts. Retinoic acid treatment reduces proliferation and tumour growth in a variety of cancers, including lung, breast, oral and skin cancer [21, 28, 29], and all-trans retinoid acid (ATRA) is currently used as treatment for acute promyelocytic leukaemia (APL) [30]. Retinoid signalling also downregulates serum response factor (SRF)-dependent genes [31], which include several cytoskeletal proteins [32], connective tissue growth factor [33] and several microRNAs [34]. While its anti-proliferative effect is well characterised, our group recently demonstrated that RAR-β activation via ATRA can regulate the mechanical activity of cancer associated fibroblasts (CAFs), including PSCs and hepatic stellate cells (HSCs) [11, 24]. These findings position RAR-β as an important player in cancer mechanobiology and an attractive target for cancer therapy. However, the mechanism by which RAR-β modulates the mechanical activity of cancer cells remains unexplored.

Here we demonstrate that the expression of RAR-β is downregulated in PDAC patients and correlates with tumour stage. We then investigate the mechanism of mechano-regulation by retinoids and identify a RAR-β binding site (RARE) on the smooth-muscle myosin light chain 2 (MLC-2) gene, leading to the downregulation of MLC-2, a key component of the contractile actomyosin machinery. Finally, we demonstrate that RAR-β-dependent MLC-2 downregulation modulates the mechanical activity of PDAC cells, including traction force generation and mechanosensing, reduces the stiffness of cancer cells, and impairs their ability to invade through the basement membrane. Together, our results shed new light into the role of retinoids as mechano-modulating drugs in pancreatic cancer.

## 2. Results

### 2.1 RAR-β expression is reduced in pancreatic ductal adenocarcinoma

The retinoic acid receptor β (RAR-β) is a tissue-specific RAR that has been postulated as a tumour suppressor [35, 36]. RAR-β expression is dysregulated or supressed in several types of cancer including lung, cervix and breast cancer [19], and the loss of RAR-β expression is associated with poor prognosis in colorectal cancer [37].

Here, we analysed the RAR-β expression status in pancreatic ductal adenocarcinoma (PDAC) and healthy patients using tissue microarrays (TMAs) and immunofluorescence staining. Healthy tissues exhibited high expression of RAR-β **(Fig. 1A and 1B)** but this expression was significantly reduced in PDAC and PDAC-adjacent tissues, with a ∼70% reduction in expression between healthy and PDAC tissue. Staining for PAN-cytokeratin, a marker of pancreatic cancer cells, revealed high expression in PDAC tissue but negligible expression in healthy tissue. Interestingly, RAR-β expression in PDAC was localised to areas of lower PAN-cytokeratin expression.

**Figure 1.**
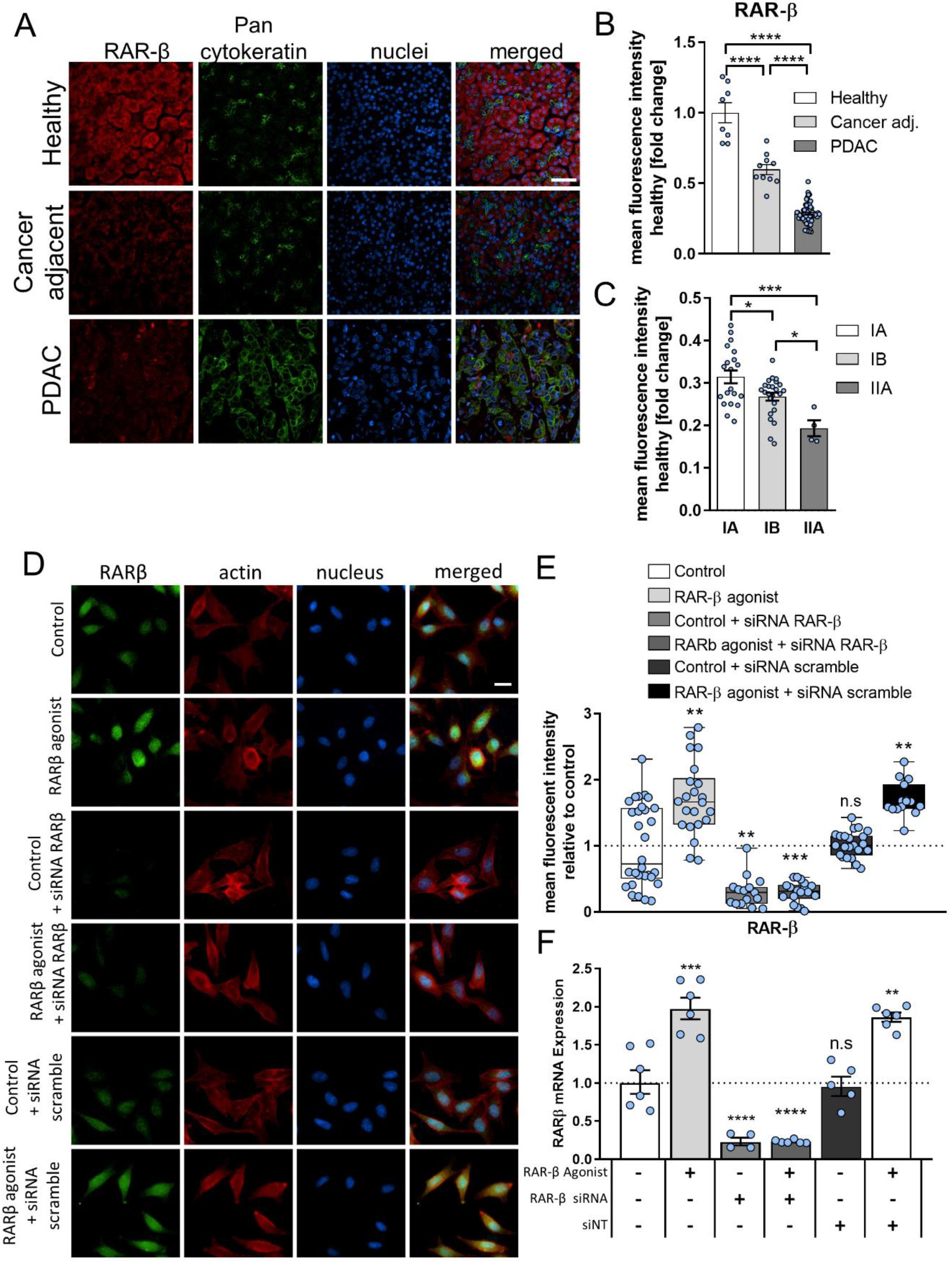
Expression of RAR-β in tissue arrays. **(A)** Representative immunofluorescence images for RAR-β (red), PAN cytokeratin (green) and DNA (blue) in healthy pancreas, cancer adjacent tissue and PDAC tissue microarrays. Scale bar represents 50 µm. **(B)** Quantification of the immunofluorescence staining in (a). Mean ± s.e.m., n=8, 10, and 60 for healthy, cancer adjacent and PDAC tissue microarrays respectively. One way ANOVA test with Tukey’s post-hoc test, **** p<0.0001. **(C)** Quantification of the mean fluorescence intensity for RAR-β on tissue micro arrays for stage IA, IB and IIA PDAC relative to healthy pancreatic tissue. Mean ± s.e.m., n= 20, 24, and 4 for stages IA, IB and IIA respectively. One way ANOVA test with Tukey’s post-hoc test, * p<0.05, *** p<0.001. **(D)** Immunofluorescence analysis of RAR-β expression at the protein level. Representative images for control, RAR-β agonist, RAR-β siRNA, RAR-β agonist + RAR-β siRNA, Non-targeting siRNA and RAR-β agonist + Non-targeting siRNA respectively. Scale bar = 20 µm. **(E)** Quantification of immunofluorescence staining in (D). Mean ± s.e.m., n=30, 22, 15, 18, 22 and 15 respectively. Markers (*) indicate significant difference relative to control by Kruskal-Wallis test with Dunn’s post-hoc test, n.s. not significant, ** p<0.01, *** p<0.001. **(F)** Relative expression of RAR-β in control, RAR-β agonist, RAR-β siRNA, RAR-β agonist + RAR-β siRNA, Scrambled siRNA (siNT) and RAR-β agonist + siNT respectively as measured by mRNA RT qPCR normalised to RPLP0. Mean ± s.e.m., n=6, 6, 4, 6, 5 and 6. Markers (*) indicate significant difference relative to control by one way ANOVA test with Dunnett’s post-hoc test, n.s. not significant, ** p<0.01, *** p<0.001, **** p<0.0001.

Analysis of cancer adjacent tissues indicated a similar trend, with increased PAN-cytokeratin expression and decreased RAR-β expression (∼50% compared to healthy tissues). Similarly, we observed a correlation between the loss of RAR-β expression and the tumour stage **(Fig. 1C)**, with a significant reduction in RAR-β expression between IA, IB and IIA tumour tissues. Together, these results suggest that the loss of RAR-β expression is associated with PDAC progression and the development of the malignant phenotype, consistent with previous findings [35, 38].

Positive RAR-β autoregulation, i.e., an increase in RAR-β expression upon retinoid treatment, has been previously reported in humans, mice and rats [39-42]. Here, we hypothesised that treatment with retinoids could restore RAR-β signalling in pancreatic cancer cells. To this end, we treated Suit-2 cells, a highly malignant PDAC cell line, with the selective RAR-β agonist CD2314 for 24 hours. Immunofluorescence analysis of RAR-β expression in Suit2 cells revealed a nearly 2-fold increase in the expression of RAR-β at the protein level upon treatment with RAR-β agonist **(Fig. 1D and 1E)**. Analysis of mRNA expression level of RAR-β via RT qPCR revealed a similar increase in RAR-β agonist-treated cells compared to control (untreated) cells **(Fig. 1F**). Knocking down of RAR-β via siRNA effectively decreased RAR-β expression at the protein and mRNA levels both in control cells and RAR-β agonist-treated cells, thus abrogating the effect of retinoids in RAR-β autoregulation. These results indicate that RAR-β autoregulation by retinoids can restore retinoid signalling in PDAC cells.

### 2.2 RAR-β regulates MLC-2 transcription

When activated by retinoids, RAR-β forms a heterodimeric (RAR-β/RXR) transcription complex, and binds to retinoid acid response elements (RAREs) on target genes to regulate their expression. Retinoids have a pleiotropic effect on a variety of cellular programmes, from proliferation and lipid metabolism [43] to the regulation of immune system [44] or the cytoskeleton [31]. Our group has previously demonstrated that retinoid treatment can induce mechanical quiescence in pancreatic and hepatic stellate cells by targeting MLC-2 [11, 24].

Myosin light chain 2 (MLC-2) is a critical regulatory component of the actomyosin machinery. Phosphorylation of MLC-2 modulates force generation by non-muscle myosin II, the primary contractile apparatus in cancer cells, and is therefore associated with the regulation of cancer cell migration, invasion and mechanosensing. MLC-2 is upregulated in different cancer types, including melanoma [45], hepatocellular carcinoma [24], oesophageal squamous cell carcinoma [46], and PDAC [47], making it an important prognosis and therapeutic target in cancer biomechanics [48]. Analysis of MLC-2 expression in normal cancer adjacent and PDAC tissue microarrays (TMAs) confirmed a 2.5-fold increase in MLC-2 expression in PDAC **(Fig. 2A and 2B)**.

**Figure 2.**
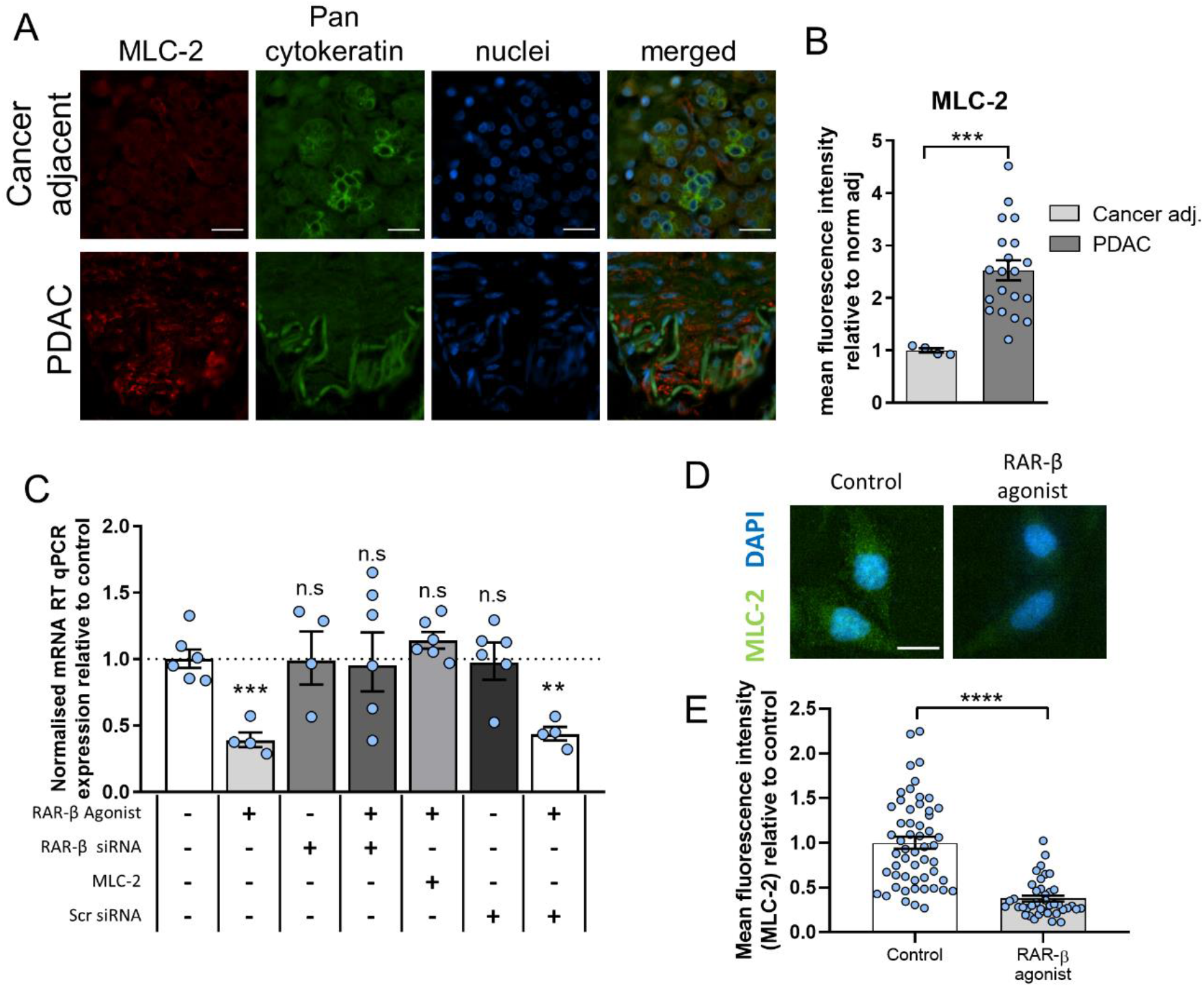
RAR-β activation downregulates expression and activity of MLC-2. **(A)** Representative immunofluorescence images for MLC-2 (red), PAN cytokeratin (green) and DAPI (blue) on cancer adjacent and PDAC tissue microarrays. Scale bar represents 50 µm. **(B)** Quantification of MLC-2 immunofluorescence staining. Mean ± s.e.m., n=4 and 20 for cancer adjacent and PDAC tissue microarrays respectively. Mann Whitney test, *** p<0.001. **(C)** Expression levels of MLC-2 quantified via RT qPCR normalised to RPLP0 and relative to control. Control, RAR-β agonist, RAR-β siRNA, RAR-β agonist + RAR-β siRNA, RAR-β agonist + MLC-2 overexpression, siNT (Scrambled) and RAR-β agonist + siNT, Mean ± s.e.m., n=6, 4, 4, 6, 6, 6 and 4 respectively. Markers (*) indicate significant difference relative to control by one way ANOVA test with Dunnett’s post-hoc test, n.s. not significant, ** p<0.01, *** p<0.001. **(D)** Representative images for MLC-2 (green) and DAPI (blue) for control and RAR-β agonist. Scale bar represents 20 µm. **(E)** Quantification of the fluorescence staining in (D). Mean ± s.e.m., n= 54 and 40 for control and RAR-β agonist respectively. Markers (*) indicate significant by Mann-Whitney test, **** p<0.0001.

To characterise the effect of retinoid signalling on MLC-2, we analysed MLC-2 expression in Suit2 cells via RT qPCR and found that treatment with RAR-β agonist for 24 hours significantly reduces the expression of MLC-2 at the mRNA level **(Fig. 2C)**. On the other hand, simultaneously treating cells with RAR-β agonist and knocking down the receptor via RAR-β siRNA inhibited the effect of the agonist and restored control levels of MLC-2 expression, indicating that the downregulation of MLC-2 expression is RAR-β dependent. Likewise, transfection with a plasmid overexpressing MLC-2 restored the expression of MLC-2 even in the presence of RAR-β agonist, whereas it had no effect on the expression of RAR-β **(Supplementary Figure S1)**. Immunofluorescence analysis of the expression MLC-2 at the protein levels revealed a similar trend **(Fig. 2D and 2E)**, with a significant reduction in MLC-2 upon treatment with RAR-β agonist. Collectively, these results indicate that retinoids transcriptionally downregulated MLC-2 expression via RAR-β.

Given the importance of MLC-2 in regulating the biomechanical activity of cancer cells, we decided to analyse the effect of RAR-β activation on the expression of YAP-1, a transcriptional effector of the Hippo signalling pathway, and a well-known marker of a mechanically active cancer cells. Using immunofluorescence, we observed that the nuclear to cytoplasmic ratio of YAP-1 (i.e., its activation) decreases upon treatment with RAR-β agonist for 24 hours **(Supplementary Figure S2)**, an effect that is abrogated when the receptor is knocked down (siRNA). These results point towards a RAR-β-dependent mechanism of mechano-modulation and prompted us to investigate the downstream effects of MLC-2 downregulation on the mechanical activity of PDAC cells.

### 2.4 RAR-β activation inhibits traction force generation, cytoskeletal stiffness and mechanosensing

MLC-2 is a fundamental regulator of actomyosin organisation and contractility, which governs the cell’s ability to generate forces and to mechanically interact with their microenvironment. A dynamic and functional actomyosin machinery is critical for cancer cells to migrate and invade other tissues, to respond to mechanical cues and to remodel their microenvironment. Based on the previous finding that RAR-β transcriptionally downregulates MLC-2, we decided to assess the effect of RAR-β activation on the mechanical activity of PDAC cells, including contractility, mechanosensing and cytoskeletal stiffness.

First, we investigated the effect of RAR-β on traction force generation using a previously established elastic micropillar platform. Micropillar arrays were fabricated from poly(dimethylsiloxane) (PDMS) via replica moulding and coated with fibronectin (FN) prior to cell seeding to enable cell attachment. Pillar displacements induced by cell-generated forces were monitored and converted to traction force maps used to quantify contractility **(Fig. 3A and 3B)**. Control Suit2 cells were able to generate a mean maximum traction force of 1.1 ± 0.1 nN (mean ± SEM, n=70 cells), comparable to other PDAC cells [49], but their contractility was significantly reduced (0.7 ± 0.1, mean ± SEM, n=53 cells, p<0.01) upon treatment with RAR-β agonist (72 hours), consistent with the downregulation of MLC-2 expression **(Fig. 3C)**. In contrast, knockdown of the receptor via RAR-β siRNA inhibited the effect of the agonist, resulting in a traction force similar to control, while overexpression of MLC-2 via transfection similarly reversed the effect of RAR-β treatment and rescued control levels of traction force generation (1.0 ± 0.1 nN, mean ± SEM, n=87 cells). These results were confirmed in a second PDAC cell line (MIA PaCa-2), with a ∼50% reduction in mean maximum force in cells treated with the RAR-β agonist CD 2314 compared to vehicle control (Supplementary Figure S3). Together these results indicate that RAR-β activation decreases cell contractility via MLC-2 downregulation, consistent with the role of the latter as a regulator of actomyosin contractility.

**Figure 3.**
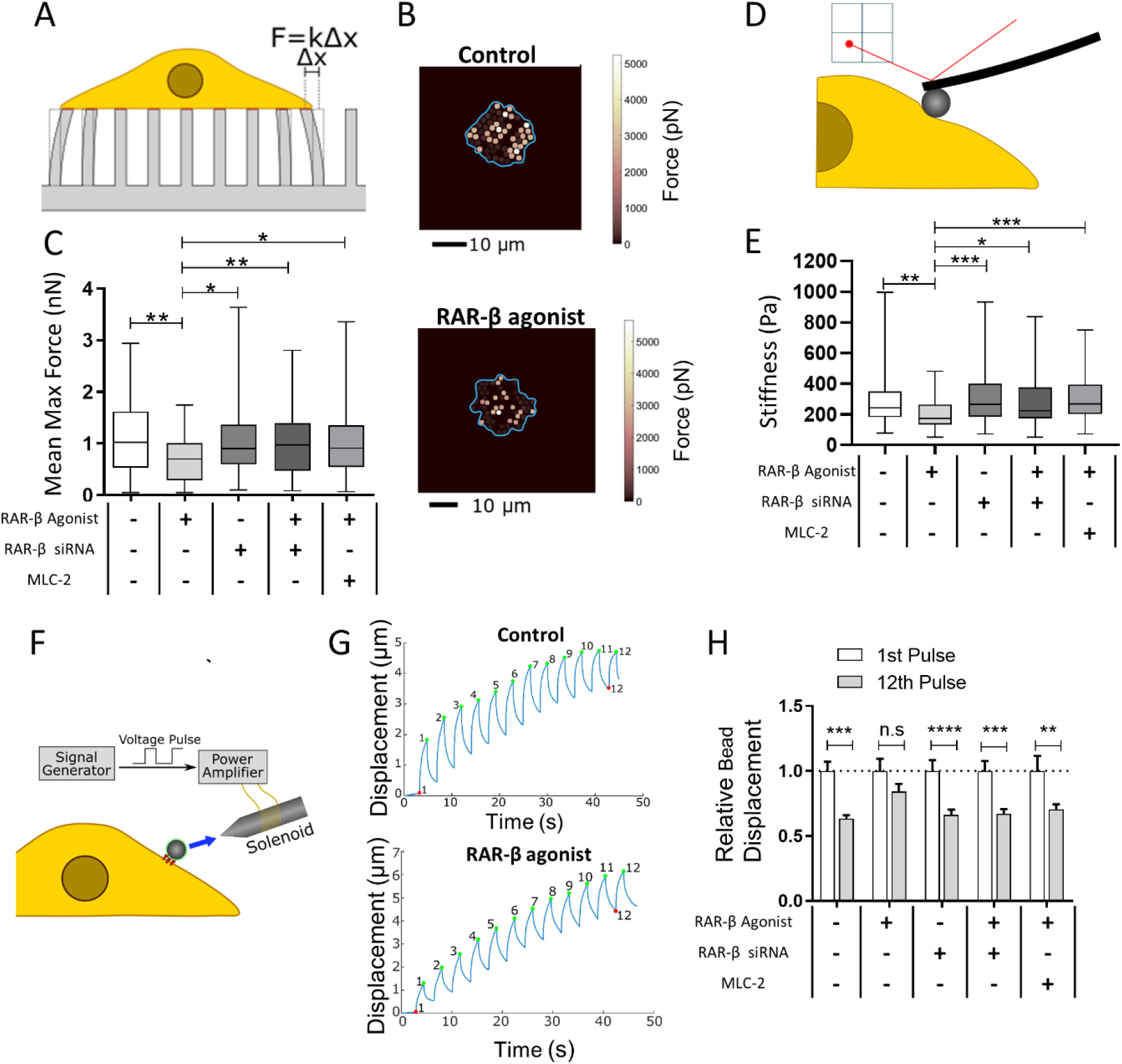
RAR-β activation impairs traction force generation, cytoskeletal stiffness and mechanosensing in pancreatic cancer cells. **(A)** Schematic of the elastic micropillar array setup to quantify cellular traction forces. **(B)** Heat map of the traction force distribution in control and RAR-β agonist-treated Suit-2 cells. The cell body is outlined in blue. Scale bar: 10 µm. **(C)** Quantification of mean maximum traction force exerted by Suit-2 cells on elastic pillars (Control, RAR-β agonist, RAR-β siRNA, RAR-β agonist + RAR-β siRNA, and RAR-β agonist + MLC-2 Overexpression, respectively). Mean ± s.e.m., n=70, 53, 97, 110 and 87 cells respectively. Kruskal-Wallis test with Dunn’s post-hoc test, * p<0.05, ** p<0.01. **(D)** Schematic of the AFM nanoindentation method to measure cell stiffness. **(E)** Cytoskeletal stiffness measured with AFM using a 15 μm bead and fitted to the Hertz model for control, RAR-β agonist, RAR-β siRNA, RAR-β agonist + RAR-β siRNA and RAR-β agonist + MLC-2 overexpression cells. Mean ± s.e.m., n=91, 58, 58, 56 cells and 64 respectively, Kruskal-Wallis test with Dunn’s post-hoc test, * p<0.05, ** p<0.01, *** p<0.001. **(F)** Schematic representation of the magnetic tweezers protocol used to measure mechanosensing in pancreatic cancer cells with the blue arrow indicating the magnetic pull applied to a fibronectin coated bead on the cell surface. **(G)** Representative bead trajectories under the pulsatile force regime (12 force pulses) in control and RAR-β agonist treated cells. A decrease in the displacement amplitude over the 12 pulses can be observed in control cells but not in RAR-β agonist treated cells. Amplitude of the 1^st^ pulse superimposed (grey) over the 12^th^ pulse (blue) for comparison. **(H)** Relative bead displacement for the 1st and 12th pulses for control, RAR-β agonist, RAR-β siRNA, RAR-β agonist + RAR-β siRNA and RAR-β agonist + MLC-2 overexpression cells respectively. A significant difference between the amplitude of the 1^st^ pulse and the 12^th^ pulse is an indicator of mechanosensing as the cell reinforces in response to the applied force. Mean ± s.e.m., n=41, 22, 28, 30 and 21 cells respectively, Wilcoxon signed-rank test, n.s. not significant, ** p<0.01, *** p<0.001, **** p<0.0001.

Cytoskeletal stiffness is another indicator of biomechanical activity that depends on the regulation of the actomyosin cytoskeleton. The ability to dynamically reorganise the cytoskeleton in response to the changing microenvironment is critical in cancer cell migration and correlates with invasive potential [50]. We analysed cytoskeletal stiffness using atomic force microscopy (AFM) to carry out nanoindentation measurements of individual cells **(Fig. 3D)**. Control Suit-2 cells showed a mean stiffness of 292 ± 20 Pa (mean ± SEM, n=91 cells), comparable to similar cell types [49], whereas cells treated with RAR-β agonist for 72 hours showed reduced stiffness (202 ± 12 Pa, mean ± SEM, n=58 cells), consistent with the decrease in MLC-2 expression **(Fig. 3E)**. We also observed that both knockdown of the RAR-β receptor (RAR-β siRNA) or MLC-2 overexpression individually recovered control-level of cytoskeletal stiffness. These results indicate that RAR-β activation causes an MLC-2 dependent reduction in cellular stiffness.

Mechanosensing is the ability for cells to sense and respond to mechanical cues. Like traction force generation, mechanosensing necessitates an intact actomyosin machinery that can dynamically reorganise in response to mechanical stimuli. Mechanical signals, including substrate stiffness, play an important role in directing cancer cell migration and invasion and are therefore one of the driving forces behind pancreatic cancer progression. Here we assessed the effect of RAR-β signalling on mechanosensing using magnetic tweezers **(Fig. 3F)**. Suit2 cells were incubated with fibronectin-coated magnetic beads, which readily attach to surface integrins, and subjected to a pulsatile force regime (12 force pulses, 6 nN, 3 seconds per pulse), while the resulting bead displacements over the 12 pulses were monitored to measure cell stiffening in response to force application.

Control Suit2 cells showed significant cytoskeletal reinforcement, with a decrease in relative bead displacement between the 1^st^ and 12^th^ pulse (∼40%, p<0.001), indicative of their mechanosensing capacity **(Fig. 3G and 3H)**. Conversely, cells treated with RAR-β agonist for 24 hours showed a decrease in cytoskeletal reinforcement, with only a ∼15% reduction in the amplitude of the 12^th^ pulse relative to the 1^st^ pulse, and a significantly larger bead displacement on the 12^th^ pulse (0.84 ± 0.06, mean ± s.e.m, n=22) compared to control cells (0.63 ± 0.02. mean ± s.e.m, n=41, p<0.001 Dunnett’s multiple comparisons test), indicative of impaired mechanosensing. Consistent with our previous findings, knockdown of RAR-β via siRNA inhibited the effect of the agonist on mechanosensing, and overexpression of MLC-2 restored the mechanosensing capacity of Suit2 cells to control levels indicating that RAR-β modulates mechanosensing in PDAC cells in an MLC-2 dependent manner.

### 2.5 RAR-β activation impairs cancer cell invasion

The first step in the metastatic journey is the breaching of the basement membrane (BM), a complex sheet-like protein bilayer that provides anchoring for the basal surface of epithelial cells and promotes apico-basal polarity. The process of epithelial-to-mesenchymal transition (EMT) that accompanies cancer progression is characterised by a loss of cell polarity, loss of cell-cell junctions and an increase in cell mobility. We recently reported that the basement membrane of PDAC differs in structure and composition from that of the healthy pancreas [49], resulting in an abnormal mechanical interaction between cancer cells and the basement membrane that promotes the breaching of the basement membrane or transmigration.

The ability to breach the initial barrier posed by the basement membrane enables tumour cells to invade neighbouring tissues and is therefore a key marker of malignancy and a critical therapeutic target in the prevention of metastasis. In order to investigate the effect of RAR-β activation on the invasive ability of cancer cells, we used a recently developed BM mimic based on mouse mesenteries [51]. Mouse mesenteries present a composition and bilayer structure similar to PDAC basement membranes and are therefore ideal models to study BM transmigration [49].

Mouse mesentery models were isolated and prepared as described by Ghose et al. [51] **(Fig. 4A)**. Suit2 cells were then cultured on the decellularised mesenteries and their transmigration across the membrane was monitored over a period of 5 days using confocal fluorescence microscopy **(Fig. 4B)**. We characterised the percentage of the cell body that penetrated the bilayer structure on days 3 and 5 **(Fig. 4C)** and found that in control Suit2 cells, the percentage of the cell body invading through the mesentery increased from 54 ± 4% at day 3 to 75 ± 3% at day 5 (mean ± s.e.m, n = 19 and 21 cells respectively). Conversely, cells treated with RAR-β agonist showed no increase in invasion between days 3 (44 ± 3%) and 5 (42 ± 2%, mean ± s.e.m, n = 21 and 18 cells respectively) and a significantly lower percentage of invasion compared to control cells at both day 3 (p<0.05) and day 5 (p<0.0001). Overexpression of MLC-2 restored the invasive potential of RAR-β agonist-treated cells, showing a high invasive potential both at day 3 (72 ± 2%) and day 5 (70 ± 2%, mean ± s.e.m, n = 22 cells). Measurement of cumulative invasion, i.e., the number of cells that fully migrated through the mesentery (per ROI) showed a similar trend **(Fig. 4D)**, with RAR-β treated cells showing decreased invasive potential over the 5-day period compared to control Suit2 cells and Suit2 cells overexpressing MLC-2.

**Figure 4.**
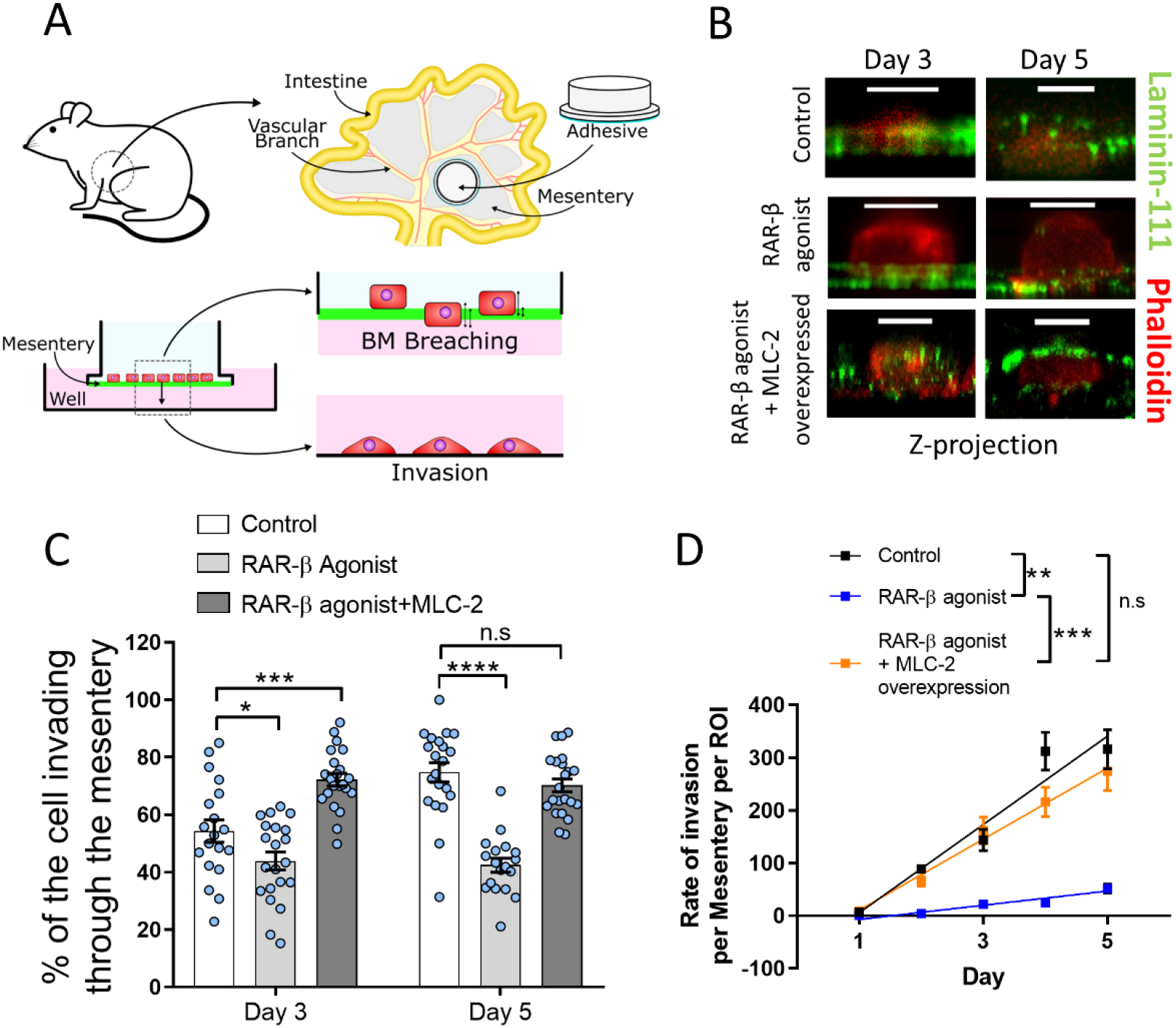
Basement membrane invasion assay. **(A)** Schematic of the mesentery preparation and invasion assay. Mesenteries from wild type mice are surgically extracted and bonded to hollow cylindrical tubes. After decellularisation, Suit2 cells are seeded on the mesentery transwells and cultured for up to 5 days. Every 24 hours, mesenteries are transferred to a new well, and the number of cells attached to the well (complete migration) are counted. Mesenteries are fixed on days 3 and 5 to quantify percentage invasion. **(B)** Confocal fluorescence images (Z-projection) of Suit2 cells (control, RAR-β agonist, and RAR-β agonist + MLC-2 overexpression) invading through mesenteries. Laminin-111 (green), actin (red). Scale bar represents 10 μm. **(C)** Quantification of the invasive capacity of Suit-2 cells. The percentage of the cell body that had invaded through the bilayer was quantified at day 3 (mean ± s.e.m., n = 19, 21 and 22 cells for control, RAR-β agonist and RAR-β agonist + MLC-2 overexpression respectively) and day 5 (mean ± s.e.m., n = 21, 18 and 22 cells for control, RAR-β agonist and RAR-β agonist + MLC-2 overexpression respectively). Markers (*) indicate significant difference relative to control by one way ANOVA test with Dunnett’s post-hoc test for Day 3 and Day 5, n.s. not significant, * p<0.05, *** p<0.001, **** p<0.0001. **(D)** Cumulative number of cells that have fully invaded through the membrane over a period of 5 days for Suit2 control, RAR-β agonist and RAR-β agonist + MLC-2 overexpression cells. Mean ± s.e.m., n= 28, 18, 26 (day 1), 53, 40, 48 (day 2), 67, 47, 56 (day 3), 71, 49, 60 (day 4), 78, 65, 68 (day 5). Lines represent linear regression model, markers (*) indicate significant difference between the slopes of the linear regression, ns not significant, ** p<0.01, *** p<0.001.

## 3. Discussion

Retinoids, the active forms of vitamin A, are a family of compounds with pleiotropic effects on cells that act through the ligand-activated transcription factors of the retinoic acid receptor (RAR) family. Loss of RAR expression, particularly the retinoic acid receptor β (RAR-β), is associated with a variety of cancers [28, 35], which prompted their use in cancer treatment, particularly for acute promyelocytic leukaemia (APL) [30]. Here, we report that RAR-β expression is reduced in pancreatic ductal adenocarcinoma (PDAC) and its downregulation correlates with tumour stage, pointing towards RAR-β as an interesting target in PDAC.

Retinoids have been previously shown to inhibit cell growth and induce apoptosis in several PDAC cell lines alone or in combination with gemcitabine [52-54]. Here we found that, when activated by retinoids, (RAR-β) downregulates the expression of Myosin Light Chain 2 (MLC-2). MLC-2 is one of the central regulatory components of myosin, in turn modulating actomyosin contractility, stress fibre formation and cortical stiffness. Force generation by actomyosin, in turn, governs the ability for cells to mechanosense their substrate, and is a key driver in mesenchymal and amoeboid migration in cancer [55] **(Fig. 5)**. These results position RAR-β as a regulator of cancer biomechanics, building on its previously established role as a cell growth inhibitor and an attractive therapeutic target in cancer.

**Figure 5.**
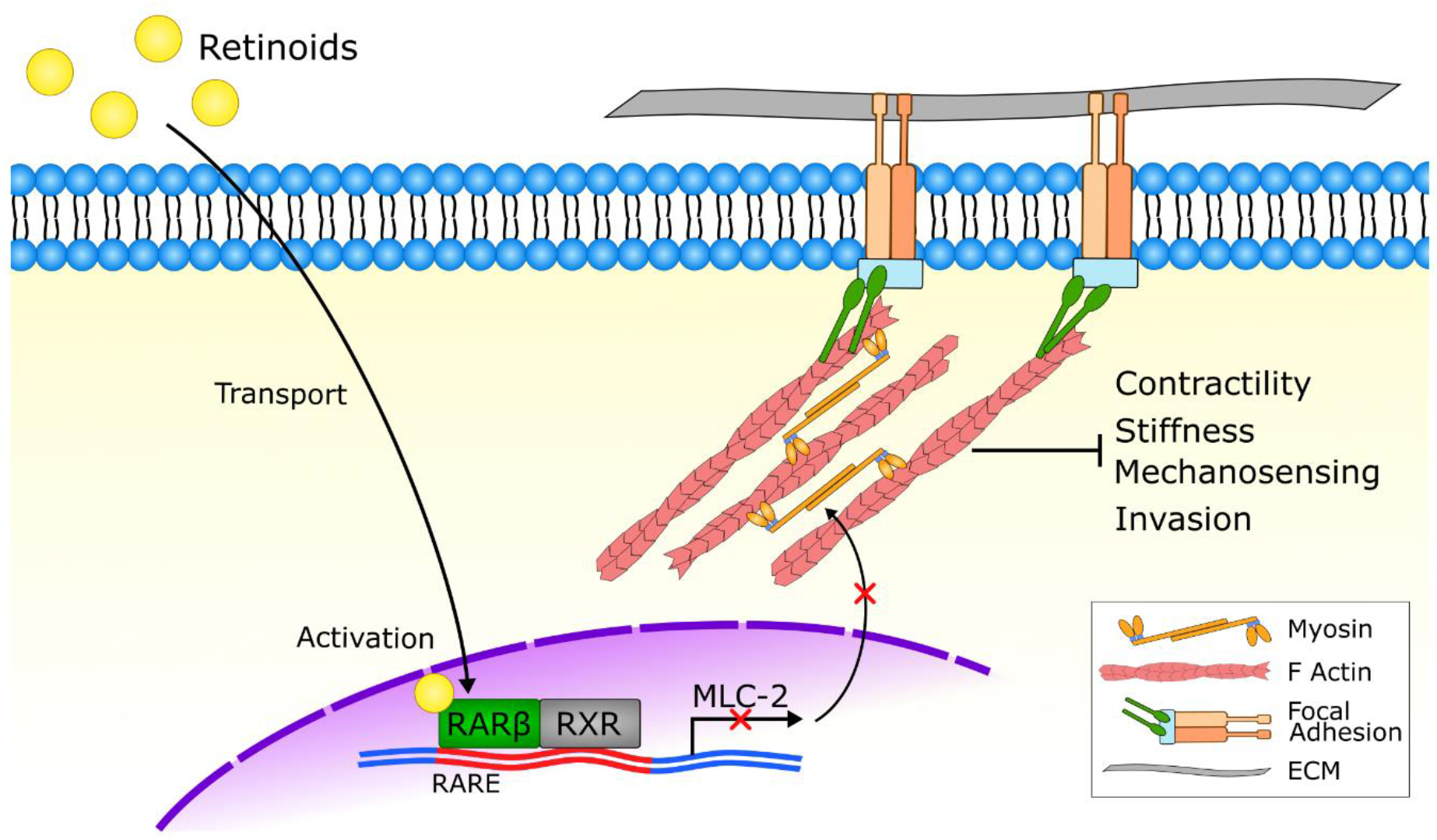
Retinoid acid receptor β modulates mechanical activity in PDAC cells via MLC-2. Retinoids activate the nuclear retinoid acid receptor β (RAR-β) along with its transcription partner (RXRs), which downregulates the expression of the myosin regulatory light chain (MLC-2), in turn decreasing actomyosin activity. By targeting the cytoskeleton, retinoids modulate traction force generation, mechanosensing, cortical stiffness and basement membrane invasion.

Despite the potential of retinoids as anticancer drugs, their clinical application has been limited by tumour chemoresistance. Resistance to retinoids has been observed in certain PDAC cell lines, as well as other types of cancer, and is often associated with a deficiency in the cellular retinoic acid-binding protein 2 (CRABP2) [52, 56]. CRABP2 stabilises retinoids in the cytosol and transports them to the nucleus, where they can bind to RARs to carry out their anti-proliferative function [57, 58]. Loss of CRABP2, or upregulation of the alternative transporter fatty acid-binding protein 5 (FABP5), results in reduced nuclear transport of retinoids and increased activation of the peroxisome proliferator-activated receptors (PPARβ/δ), which has an anti-apoptotic effect. Interestingly, the relative expression of these retinoids transporters has been proposed as a marker of retinoid sensitivity and could be used to stratify PDAC patients according to their responsiveness to retinoids [59, 60].

The loss of RAR-β expression has also been proposed as a mechanism of chemoresistance, which is often attributed to epigenetic (methylation) changes in the RAR-β promoter, leading to its silencing [52, 56]. The positive autoregulation by RAR-β that we have observed in PDAC cells could be essential to the success of retinoids as mechano-modulating drugs, increasing PDAC sensitivity to retinoid treatment. Future work will be required to investigate the mechanisms of transport and bioavailability of retinoids in order to design effective therapeutic strategies.

In recent years, the role of mechanical cues, such as tissue stiffness, in the progression of cancer has come into focus [61, 62], prompting the development of novel therapies that modulate or reprogram the biomechanical activity of both cancer cells and cancer associated fibroblasts (CAFs)—such as PSCs. Here we have shown that RAR-β can modulate the biomechanical activity of cancer cells via transcriptional repression of MLC-2. The decrease in contractility, impaired mechanosensing and the reduction in YAP nuclear localisation are all indicative of mechanical quiescence, highlighting the ability for retinoids to address the mechanical drivers of cancer.

Mechanically activated PSCs develop and maintain the aberrant microenvironment that drives PDAC progression, characterised by excessive extracellular matrix (ECM) deposition, remodelling and stiffness. Anti-stromal therapies that aim to deplete the tumour ECM, however, have proven to be unsuccessful, in some instances resulting in more aggressive tumours [63, 64]. The role of the stroma in tumour progression is complex and changes along cancer’s spatiotemporal evolution. Consequently, research efforts have shifted towards anti-stromal strategies that impair the crosstalk between cancer cells and CAFs [65-67]. Our group and others have previously shown that retinoids can mechanically reprogram PSCs by decreasing endogenous force generation, reducing mechanosensing and inhibiting matrix remodelling [11, 68-70]. Retinoid treatment could therefore act synergistically to modulate the mechanical activity of both cancer cells and CAFs, directly targeting the mechanical interaction between cancer cells and their microenvironment.

Here we have also observed that retinoid treatment decreases the invasive potential of PDAC cells, pointing towards a potential role in the prevention of metastasis. Metastasis is a complex, multi-stage process that involves cancer cell migration, intravasation and colonisation of distant tissues, all of which rely on rapid actomyosin reorganisation [71]. Cell contractility has also been shown to regulate the secretion of matrix metalloproteinases (MPPs) [12], which are required to break down and remodel the ECM. Moreover, retinoids can regulate the mechanical activity of HSCs, the primary population of liver fibroblasts responsible for liver fibrosis. As the liver is the primary metastatic site for PDAC [24, 72], retinoids could prevent the formation of a pre-metastatic niche. These findings suggest that targeting the RAR-β/MLC-2 axis could be a compelling strategy to impair cancer metastasis, although future studies using animal models will be required to elucidate the systemic effects of retinoids and to understand the interaction between their mechano-modulating and their anti-proliferative effects.

## 4. Materials & Methods

### 4.1 Cell culture and treatment

Suit2-007 cells (S2, passages 5-8) were cultured in Dulbecco’s Modified Eagle’s Media (Sigma-Aldrich, D8437) supplemented with 10% v/v FBS (Cat No.F7524, Sigma Aldrich, Dorset, UK), 2 mM L-glutamine (Cat No.G7513, Sigma Aldrich, Dorset, UK), 1% v/v penicillin/streptomycin (Sigma-Aldrich, P4333) and 1 % v/v fungizone Amphotericin B (Gibco, UK, 15290-026). Cells were incubated at 37°C, with 5% CO_2_. Media was changed every 3-4 days, and cells were subcultured at 70-80% confluency. For all RAR-β agonist treatments, cells were exposed to 1 μM RAR-β agonist (Tocris, CD 2314) 24h prior to experiments. Gene transfection was performed 48h prior to experiments, utilising the Neon transfection system (Thermo Fisher Scientific) with 2 μg MLC-2 plasmid (pEGFP-MRLC1, Addgene, #35680), 10 μg RAR-β siRNA (Santa Cruz, SC-29466) or 10 μg control siRNA (siRNA-Scr, Santa Cruz, SC-37007).

### 4.2 Antibodies and Immunostaining

Suit2 cells were seeded on coverslips previously treated with Fibronectin (Gibco, PHE0023) for 45 minutes and washed with D-PBS (Sigma-Aldrich, D8537). Cells were fixed with 4% paraformaldehyde (Sigma-Aldrich, P6148) diluted in PBS for 10 min. Fixed cells were permeabilised with 0.5% saponin (Sigma-Aldrich, 47036) diluted in PBS for 10 min. Cells were then incubated in a blocking solution of 1% bovine serum albumin (BSA; Sigma-Aldrich, A8022, Lot SLBT9577) and 22.52 mg/mL UltraPure Glycine (Invitrogen, Lot 18C2856902) diluted in PBS with 0.1% Tween 20 (Sigma-Aldrich, P1379) for 30 min. Cells on coverslips were incubated with diluted primary antibodies in PBST against myosin light chain-2 (MLC-2, Cell Signaling Technology, 3672S, Rabbit, 1:100), RAR-β (Abcam, ab53161, Rabbit, 1:50) and YAP (Cell Signaling Technology 4912S, Rabbit, 1:50) at 4 °C overnight. This was followed by incubation with goat anti-rabbit 488 (Life Technologies, A11034, 1:200) secondary antibody and Phalloidin-594 (Abcam, ab176757, 1:200) for 1 hour at room temperature. Cells were then mounted with Prolong Gold antifade reagent containing DAPI (ThermoFisher Scientific) and imaged on a Nikon Eclipse Ti-E microscope with a 40× objective (Nikon, Kingston-upon-Thames UK).

Fixed Suit2 cells on mesenteries were permeabilised using 0.5% Triton X-100 (Sigma-Aldrich, T8787). Fixed samples were blocked using 2% BSA. PBS was used to block non-specific binding for 30 min. Samples were then incubated with primary antibodies for laminin (Rabbit, Sigma-Aldrich, L9393, 1:100) at room temperature for 1 h. This was followed by incubation with goat anti-rabbit Alexa Fluor 488 (ThermoFisher, A11034, 1:200) secondary antibody and Phalloidin-594 (Abcam, Ab176757, 1:200 dilution) for 1 hour at room temperature. Samples were then mounted using Prolong Gold antifade reagent with DAPI (Thermo Fisher Scientific) and imaged using confocal microscopy with a Nikon Eclipse Ti-E microscope, 60× objective (Nikon, Kingston-upon-Thames UK). 3D confocal images were acquired with 0.25 μm z-section spacing.

### 4.3 Immunofluorescence Image Analysis

Widefield fluorescent images were taken with a Nikon Ti-e Inverted Microscope (Ti Eclipse, C-LHGFI HG Lamp, CFI Plan Fluor 40 × NA 0.6 air objective; Nikon Europe, Amsterdam, Netherlands; Neo sCMOS camera; Andor, Belfast, UK) with NIS elements AR software. Staining fluorescence intensity was quantified in Fiji [73] using the “mean gray value” parameter applied to a region of interest (ROI). Regions of interest (ROI) for individual cells were segmented based on their actin staining (red channel). For YAP nuclear localisation, nuclear ROIs were defined through automated thresholding of the DAPI (nuclear) channel, and cytoplasmic ROIs were defined as the whole cell ROI with subtracted nuclear ROI. Ratios of the nuclear to cytoplasm fluorescence intensities (“mean gray value”) were calculated in order to analyze the localization of YAP in the different cells. Unless otherwise specified, individual data points represent the average intensity (or average nuclear/cytoplasmic intensity ratio) for each field of view (10-20 cells quantified per field of view) across three independent replicates.

### 4.4 Tissue microarray staining

For quantification of the levels of RAR-β and MLC-2 in human pancreatic tissues from healthy, cancer adjacent and PDAC tissues, tissue micro arrays (TMAs) were obtained from Biomax (Catalogue number PA803 for RAR-β, PA242e for MLC-2). Formalin fixed, paraffin embedded pancreatic ductal adenocarcinoma and normal tissue arrays (US Biomax inc., cat. PA803) were dewaxed in histoclear (National Diagnostics, cat. HS-200) and rehydrated in decreasing concentrations of ethanol. Subsequently, samples underwent heat induced epitope retrieval in pH 6.0 citrate buffer for 30 minutes at 95°C. After cooling to room temperature, the array was washed in tris-buffered saline (TBS) plus 0.025% Triton X-100 (Sigma, T8787) and blocked in 10% normal goat serum (Sigma-Aldrich) with 1% BSA in TBS for 2 h at room temperature. Blocked samples were incubated overnight at 4°C with anti-RAR-β antibodies (Abcam, ab53161, Rabbit, 1:100) or anti-MLC-2 antibodies (Cell Signaling Technology, 3672S, Rabbit, 1:100) and anti-PAN cytokeratin (Abcam, ab6401, 1:250) in TBS with 1% BSA. Following the incubation, samples were washed in TBS plus 0.025% Triton X-100 with gentle agitation and incubated with secondary antibodies, goat anti-mouse Alexa Fluor 488 (ThermoFisher, A-11030) and goat anti-rabbit Alexa Fluor 546 (ThermoFisher, A-11035) diluted 1:400 in TBS with 1% BSA for 1 hour at room temperature. Tissue microarrays were mounted using Prolong Gold antifade reagent with DAPI (ThermoFisher Scientific) and imaged with a Nikon Eclipse Ti-E microscope, 20x objective (Nikon, Kingston-upon-Thames UK). Mean fluorescence intensity of the images was quantified using Fiji [73].

### 4.5 Real-time qPCR

Total RNA was extracted using the RNeasy Mini kit (Qiagen, 74104) and 1 µg of total RNA was reverse-transcribed using the High-Capacity RNA-to-cDNA kit (Applied Biosystems, 4387406) according to the manufacturer’s instructions. Real-time quantitative PCR (qPCR) was performed using the SYBR Green PCR Master Mix (Applied Biosystems, 4309155). Primer sequences: RPLP0: forward 5’-CGGTTTCTGATTGGCTAC-3’ and reverse 5’-ACGATGTCACTTCCACG-3’; RAR-β: forward 5’-AATAAAGTCACCAGGAATCG -3’ and reverse, 5’-CAGATTCTTTGGACATTCCC -3’; MLC-2: forward, 5′-ATCCACCTCCATCTTCTT-3′ and reverse, 5′-AATACACGACCTCCTGTT-3′. RT qPCR data was analysed using the 2^-ΔΔCT^ method with RPLP0 as endogenous gene control. Error bars for RT qPCR data were calculated as 2^-(ΔΔCT±SEM)^ as described in [74].

### 4.6 Magnetic tweezers

Magnetic beads (4.5 µm, Dynabeads M-450, Thermo Fisher Scientific) were coated with fibronectin (Gibco, PHE0023) following the manufacturer’s instructions. Suit2 cells were incubated with these fibronectin-coated beads for 30 minutes at 37 °C and then thoroughly washed with PBS to remove unbound beads. Individual beads were then subjected to a pulsatile force regime consisting of a 3 s, 6 nN pulse of force, followed by a 4 s period of rest, repeated for 12 total pulses over a 100 s time course. The bead trajectories were recorded in a Nikon Ti-E inverted microscope (Ti-Eclipse, C-LHGFI HG Lamp, CFI Plan Fluor 40× NA 0.6 air objective; Nikon; Neo sCMOS camera; Andor) with NIS elements AR software and analysed using a custom MATLAB script. The amplitudes of each pulse were extracted from bead trajectories and normalised to the 1^st^ pulse. The amplitudes of the 1^st^ and 12^th^ pulse were compared to quantify the decrease in amplitude of the bead movement as a result of cytoskeletal reinforcement.

### 4.7 Elastic micropillar arrays

Elastic micropillar arrays were fabricated in polydimethylsiloxane (PDMS). PDMS (Sylgard 184, Dow) was mixed in a 1:10 weight ratio according to the manufacturer specifications, poured on a silicon mould and cured at 60°C for 1 hour, resulting in PDMS with a spring constant of k=1.36 nN/µm. After curing, PDMS pillars were peeled-off in PBS and stored at 4°C. Prior to seeding cells, PDMs pillars were coated with fibronectin (FN) (10 μl/mL in PBS) for 1 hour at 37°C. Cells were seeded on the FN-coated pillars and incubated for 1 hour at 37°C and 5% CO_2_ before analysing them. Each sample was analysed at 37°C for a maximum of 30 minutes in a Nikon Ti-e Inverted Microscope (Ti Eclipse, C-LHGFI HG Lamp, CFI Plan Fluor 40X NA 0.6 air objective; Nikon; Neo sCMOS camera; Andor) with NIS elements AR software. Each cell was recorded for 1 minute with a frame rate of 1 frame per second. Data analysis was carried out with a custom MATLAB script to quantify pillar deflection and traction forces exerted on each pillar were calculated based on the deflection of the pillar and the spring constant.

### 4.8 Cell stiffness

Cell stiffness was analysed with AFM nanoindentation using a Bruker Nanowizard 4 in contact – force spectroscopy mode. Nanoindentation measurements were carried out with an MLCT silicon nitride probe (Buker) with a nominal spring constant of 0.03 nN/m with a 15 µm polystyrene bead attached to the tip. Prior to cell analysis, the sensitivity of the probe was calibrated by measuring the force– distance slope in the AFM software on an empty petri dish region. Nanoindentation of individual cells was conducted at 5 µm/s to a set point of 1 V (∼1nN force set point). Cells were indented at a point between the nucleus and the cell periphery to characterise the cytoskeletal stiffness. The stiffness (Young’s modulus) of individual cells was calculated from the force-distance curves using the AFM software and the Hertz model [75].

### 4.9 Mesentery isolation and invasion assay

Mesenteries were isolated and prepared as described in [51]. 1.5 mL Eppendorf tubes were cut to create a tube of constant diameter approximately 1 cm in height to be used as a frame for the mesenteries. Mesenteries were isolated from mice intestines (generously provided by Dr Charlotte Dean from Imperial College London) using 3M Vetbond tissue adhesive (Vetbond, 3M-1469sb). Mesenteries were immediately incubated for 1 h in sodium azide (NaN_3_, Sigma-Aldrich, S2002, lot BCBL9058V) diluted in phosphate buffered saline prepared in our laboratory (PBS) for preservation, and decellularised by 1 h incubation with 1M ammonium hydroxide solution (NH_4_OH, Sigma-Aldrich, 09859, lot BVBV8688). Mesenteries were then washed and stored in Dulbecco’s PBS (D-PBS; Sigma-Aldrich, D8537) at 4°C. Cells were collected using 0.25% of trypsin-EDTA solution (Sigma-Aldrich, T4049), centrifuged, resuspended in serum-free media, and seeded inside the Eppendorf ring on top of mesenteries that were previously placed in wells with serum containing (10% FBS) media to drive the invasion assay. Mesenteries were transferred to a new culture dish every 24 h.

Mesenteries were fixed on days 3 and 5 with 4% paraformaldehyde for 10 min. Invasion was assessed using confocal fluorescence microscopy (Ti Eclipse, Nikon). For each mesentery, an average of 5 randomly selected fields of view were analysed, with an average of 10 cells per field of view. Percentage cell invasion was quantified from cross-sectional confocal images as the ratio of the height of the cell below the mesentery bilayer to the total cell height. For cumulative invasion, after the mesenteries were transferred to a new well, the number of cells that had invaded thought the mesentery and attached to the bottom of the wells were quantified within randomly selected regions of interest (ROI) by imaging on a bright field inverted microscope (Motic, AE31 trinocular). The number of cells per mesentery per ROI were quantified every 24 hours, and cumulated over a period of 5 days.

### 4.10 Statistical Analysis

All statistical analyses were conducted with the Prism software (version 8, GraphPad). Data were collected from multiple repeats of different biological experiments to obtain the mean values and s.e.m. displayed throughout. P values have been obtained using Mann-Whitney on unpaired samples with parametric tests (t-test) used for data with a normal distribution (following Normality test). ANOVA with post hoc Tukey’s or Dunnett’s test were used to perform multiple comparison test on normally distributed data. Kruskall-Wallis test followed by post hoc Dunn’s multiple comparison test was used for multiple comparison tests with data that does not follow a normal distribution. Significance was set at P < 0.05 where graphs show significance through symbols (*0.05 < P < 0.01; **0.01 < P < 0.001; ***0.001 < P < 0.0001; ****P < 0.0001).

## 5. Conclusion

Retinoids are a family of compounds derived from vitamin A with important functions in cell differentiation and homeostasis which have been explored in a clinical context as a treatment for cancer. Here we demonstrate that targeting the retinoid acid receptor RAR-β can modulate the biomechanical activity of PDAC cells via the transcriptional downregulation of MLC-2, a critical regulator of the actomyosin cytoskeleton. Using a suite of biomechanical characterisation tools, we demonstrate that RAR-β treatment results in reduced contractility, stiffness, mechanosensing and invasiveness in PDAC cells. Our work positions the RAR-β/MLC-2 axis as an important player in therapies aimed at mechanically modulating the tumour and its microenvironment.

## Supplementary Material

**Supplementary Figure S1.**
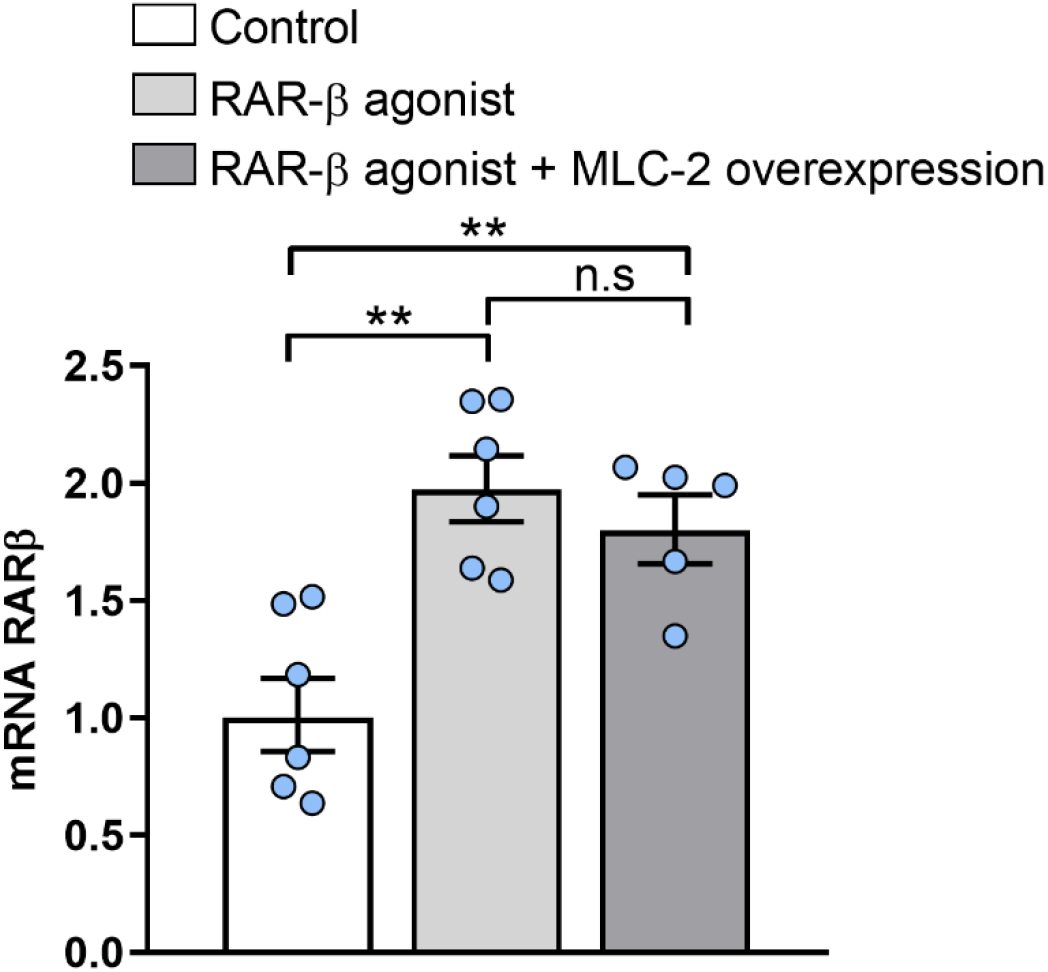
MLC-2 overexpression does not affect RAR-β expression. Relative expression of RAR-β in Control, RAR-β agonist, RAR-β agonist + MLC-2 overexpression respectively as measured by mRNA RT qPCR normalised to RPLP0. Control and RAR-β agonist data reproduced from Figure 1F. Mean ± s.e.m, n=6, 6, 5. Markers (*) indicate significant difference by one way ANOVA test with Tukey’s post-hoc test, n.s not significant, ** p<0.01.

**Supplementary Figure S2.**
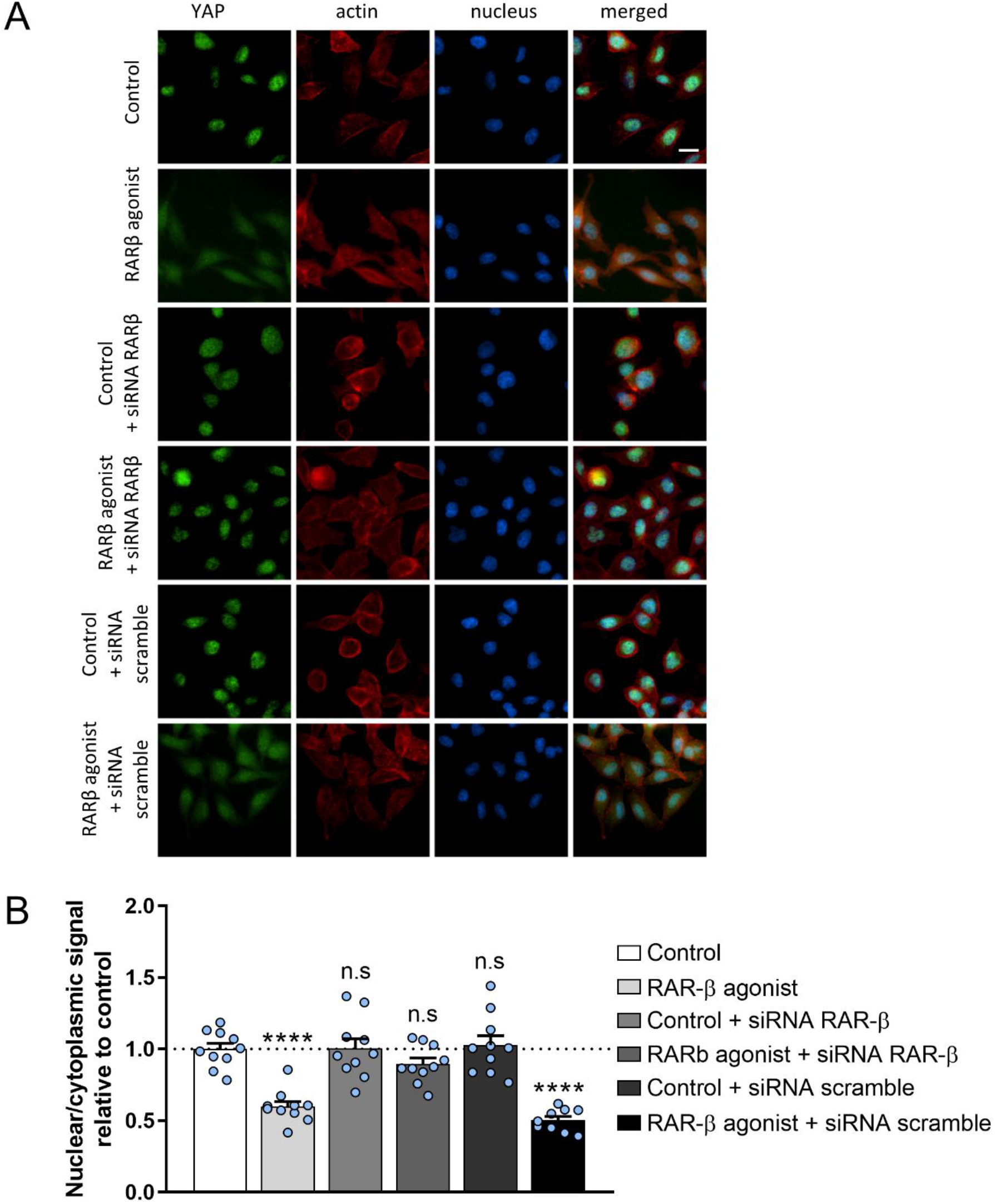
RAR-β activation induces mechanical quiescence in Suit-2 cells. **(A)** Representative images of immunofluorescence analysis of YAP-1 localisation within the cell for control, RAR-β agonist, RAR-β siRNA, RAR-β agonist + RAR-β siRNA, and Scramble siRNA and RAR-β agonist + Scramble siRNA. YAP (green), actin (red) DAPI (blue). Scale bar = 20 µm. **(B)** Quantification of the nuclear/cytoplasmic ratio of YAP-1 location based on the staining in (A). Mean ± s.e.m., n=10 fields of view for control, RAR-β agonist, RAR-β siRNA, RAR-β agonist + RAR-β siRNA, and Scramble siRNA, n=9 fields of view for RAR-β agonist + Scramble siRNA respectively. Markers (*) indicate significant difference relative to control by one way ANOVA test with Dunnett’s post-hoc test, n.s. not significant, ** p<0.01, *** p<0.001.

**Supplementary Figure S3.**
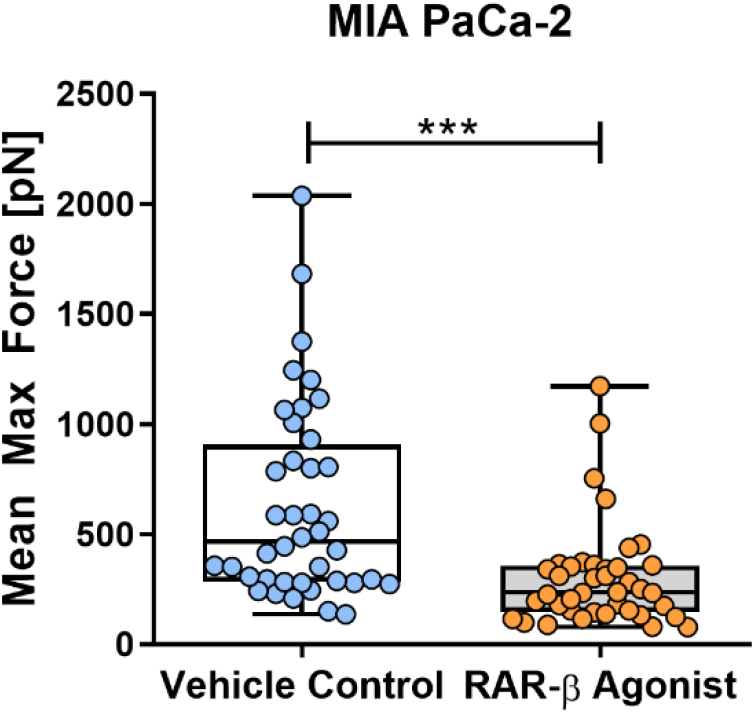
RAR-β regulates MLC-2 expression and traction force generation in MIA PaCa-2 cells. Quantification of mean maximum traction force exerted by MIA PaCa-2 cells on elastic pillars. n=40 cells for vehicle control (DMSO) and RAR-β agonist (CD 2314) treated cells. Symbols (*) denote significant difference by Mann-Whitney test, *** p<0.001.

